# Microbial adaptation to venom is common in snakes and spiders

**DOI:** 10.1101/348433

**Authors:** E. Esmaeilishirazifard, L. Usher, C. Trim, H. Denise, V. Sangal, G.H. Tyson, A. Barlow, K.F. Redway, J.D. Taylor, M. Kremyda-Vlachou, S. Davies, T. D. Loftus, M.M.G. Lock, K. Wright, A. Dalby, L.A.S. Snyder, W. Wuster, S. Trim, S.A. Moschos

## Abstract

Animal venoms are considered sterile sources of antimicrobial compounds with strong membrane disrupting activity against multi-drug resistant bacteria. However, bite wound infections are common in developing nations. Investigating the oral and venom microbiome of five snake and two spider species, we evidence viable microorganisms potentially unique to venom for black-necked spitting cobras (*Naja nigricollis*). Among these are two venom-resistant novel sequence types of *Enterococcus faecalis*; the genome sequence data of these isolates feature an additional 45 genes, nearly half of which improve membrane integrity. Our findings challenge the dogma of venom sterility and indicate an increased primary infection risk in the clinical management of venomous animal bite wounds.

**One Sentence Summary:** Independent bacterial colonization of cobra venom drives acquisition of genes antagonistic to venom antimicrobial peptides.

## Main Text

The rise of multi-drug resistant (MDR) bacterial infections suggests the end of the antibiotic golden era might be approaching fast. Discovery of novel antimicrobials is therefore an urgent priority of exceptional socioeconomic value. Crude preparations of animal venoms exhibit strong antibiotic potencies, including against clinical MDR bacterial isolates such as *Mycobacterium tuberculosis* (1). With antimicrobial properties described for crotalid (pit viper) venom as early as 1948 (2), relevant compounds have been isolated from most animal venoms, including those of spiders, scorpions, and insects, as well as aquatic species. Examples include phospholipase-A2 enzymes, L-amino acid oxidases, cathelicidins, C-type lectins, and hydrophobic/cationic peptides, as well as venom toxin domains (3), which may act by physically disrupting bacterial cell membranes through pore formation (4, 5). Accordingly, venomous animal bite or sting (envenomation) wound infections are considered rare (6) and are attributed to secondary infection (7). Yet over three quarters of snake bite victims may develop mono- or polymicrobial envenomation wound infections, characterized by *Bacteroides*, *Morganella, Proteus*, *and Enterococcus* (8, 9) – bacterial taxa commonly found in the gut. Indeed, *Enterococcus faecalis and Morganella morganii* have been independently reported as the most common Gram positive and Gram negative envenomation wound infections across several countries (8–10). Historically associated with the oral snake microbiome (11), these bacteria are thought to originate from prey faeces (12) persisting in the snake oral cavity (10) with a diversity similar to that of the snake gut (13). Yet no ‘fixed’ oral microbiome was observed in early systematic studies, beyond a seasonal variation of diversity (14). Curiously, non-venomous snake mouths were reportedly more sterile than those of venomous snakes (14), a counterintuitive finding independently reproduced elsewhere (10). More recently, the oral microbiome of the non-venomous Burmese python (*Python bivittatus*) has also been reported to be native and not derived from prey guts (15).

As venom glands are connected to the tip of envenomation apparatus via a persistently open duct which is continuously exposed to the environment (16), envenomation apparatus could be compared to clinical catheterisation assemblies: a transcutaneous needle resting on a non-sterile environment, connected to a continually open duct, leading to a liquid vessel. Such devices develop biofilms within a few days, making weekly catheter replacement necessary (17). Unlike the high flow rates of catheters, however, envenomation apparatus normally ejects venom only sporadically. Captive snakes are often fed weekly and can fast for months whereas large arachnids are fed typically on a monthly basis. Wild animals may also undergo hibernation for several months when venom expulsion frequency can be assumed to be zero. We therefore postulated that the anatomy of envenomation apparatus allows it to be colonised by microbes and that their intermittent use may facilitate bacterial persistence, adaptation and establishment within antimicrobial venom.

## Results

### The snake venom microbiome varies on account of host species and not on account of the oral flora

Applying established culture-free methods (18) on commercially available *Bothrops atrox* venom (*fer-de-lance*; viperidae) and a venom sample from a captive *Bitis arietans* (African puff adder), we first optimised microbial DNA extraction for this unusual biological matrix **(Fig. S1)**. Given animal availability, behavioural, and sampling limitations, we next focused our efforts on five snake, two spider and two scorpion species (**Table 1**). We collected a swab (O) of the oral cavity (snakes) or fang/aculeus surface (spiders/scorpions) and two consecutive envenomation samples (E1 and E2), expecting the second venom sample to have fewer contaminants from bacterial plugs possibly forming on the envenomation apparatus. In agreement with previous reports (11, 14), principle coordinate analysis and unsupervised clustering (**Fig. S2**) failed to discriminate the swab microbiomes by host species, suggesting the common diets and water sources in captivity would have the biggest impact on the swab data.

**Table 1:**
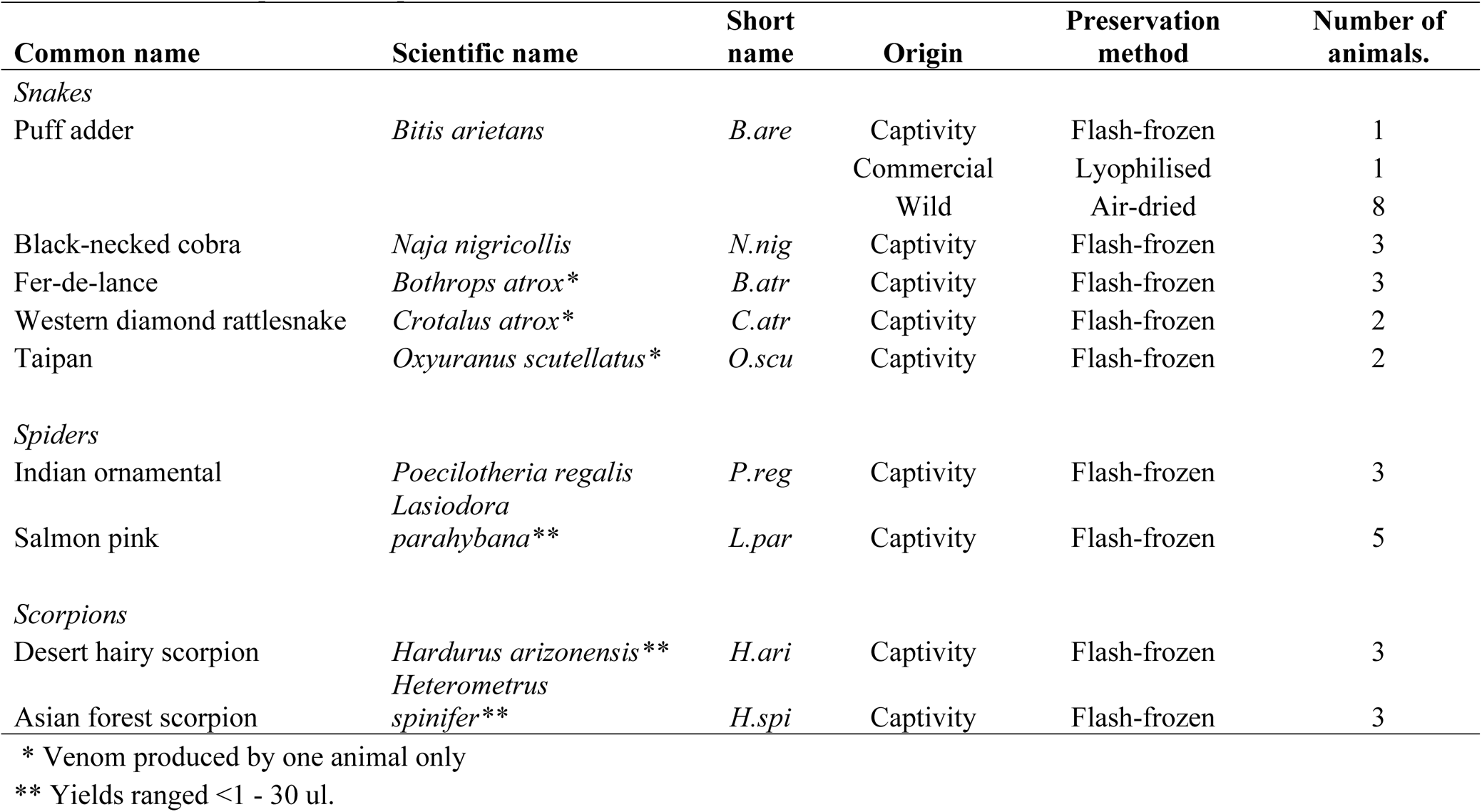
Animals sampled for the presence of microbiomes in venom.

This led us to hypothesise that captive animals would feature more closely related microbiota in their venoms compared to commercial or wild samples. We therefore compared the venom microbiomes of all snakes using the same approach (**Fig. 1**). High Shannon-Weiner indices indicated considerable diversity in snake venom microbiomes, however, surprisingly closer relationships were observed between *B. arietans* and other *Viperidae*, despite samples spanning captive and wild animals; an exception was *B. atrox* venom, which was characterized principally by *Gammaproteobacteria.* Focusing on *B. arietans* also failed to cluster samples by origin (**Fig. S2**), despite the disparate locations across South Africa where wild *B. arietans* samples were collected (**Fig. 1D**). In contrast, *N. nigricollis* microbiomes largely formed a distinct cluster (**Fig. 1**) characterized by *bacteroidia* (*bacteroidaceae*), a taxon less common among *Viperidae*. This could reflect anatomical differences in elapid (cobra) fang location at the front of the mouth compared to the sheathed nature of the longer, hinged viperid fangs, whose tips rest at the back of the oral cavity. In contrast, spider species did not seem to influence venom microbiome consistency and exhibited lower biodiversity (**Fig. S4)**. These results likely reflected vertebrate/invertebrate anatomical differences and the limited venom yield from invertebrates (<1-30 μl) vs snakes (100-1,000 μl).

**Figure 1:**
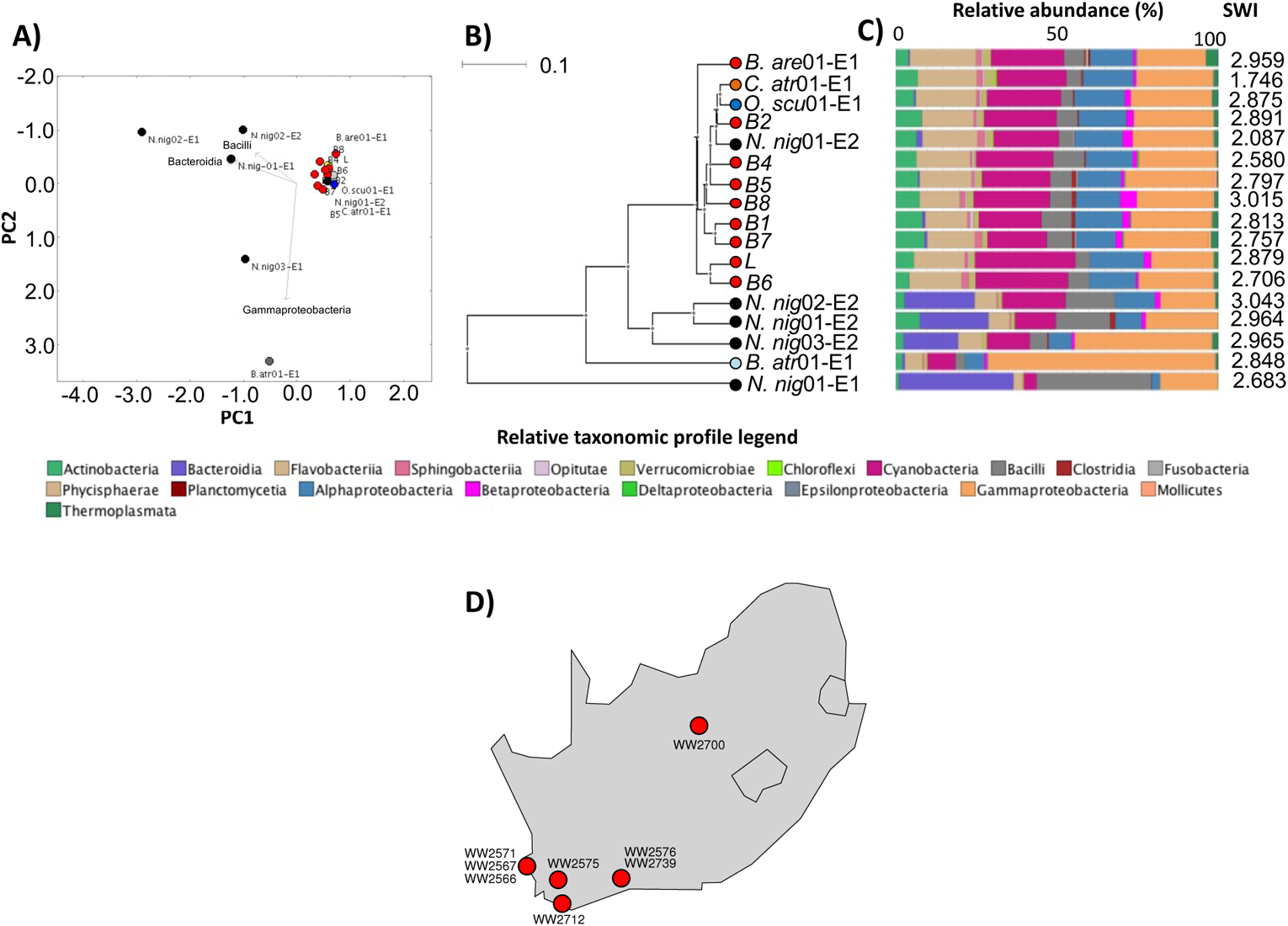
Snake venom microbiomes cluster on account of host species. Viperid venom microbiomes cluster separately from *N. nigricollis* with the exception of *B. atrox* as determined by A) PCoA, B) UPGMA tree and C) class-level taxonomic profiling following 16S rRNA phylogenetic analysis. Dots in (A) and (B) are coloured by species (red: *B. arietans*; black: *N. nigricollis*; light blue: *B. atrox*; orange: *C. atrox;* dark blue: *O. scutallatus*), represent data of captivity individuals, are labelled with short species name, enumerated for individual number and identified for the envenomation number (E1 or E2) of the sample. The 8 wild (red dots B1-B8) and the commercially sourced, lyophilised (red L dot) *B. arietans* sample are independently labelled. Relative taxonomic diversity profiles in (C) are aligned to the UPGMA tree sample labels, with the Shannon-Weiner Index (SWI) of each sample indicated. The geographical origin of the wild *B. areitans* samples collected in South Africa are shown in D.

### A fifth of the *N. nigricollis* venom microbiome is distinct to that of fangs

Encouraged by the distinctive bacterial taxonomies in *N. nigricollis* venom, the availability of animals under controlled conditions, and the paired nature of the fang swab and envenomation samples, we delved deeper into this dataset. The fang microbiomes appeared to form a distinct cluster to that of venom microbiomes (**Fig. 2A, B)** suggesting the venom gland might be a distinct ecological niche (**Fig. 2C**). We therefore asked if any bacterial taxa were unique to subsets of these samples. Operational taxonomic unit (OTU) incidence analysis within each animal (**Fig. 2D**) suggested some 60% of OTUs were shared between corresponding venoms and fangs. Yet, importantly, up to 20% of these appeared to be unique to venom, and some 15% were unique to the fang (**Fig. 2E**), indicating an OTU continuum between the two microenvironments, with unique taxa in each site. Common sample types also featured a majority of common taxa, and OTUs unique to each site in each animal **(Fig. 2F**). However, taxa unique to each sample type (O, E1 or E2) were rarely found across all animals. These results suggested that although the microbiome between each snake fang and venom was largely common, venom contained distinct organisms.

**Figure 2:**
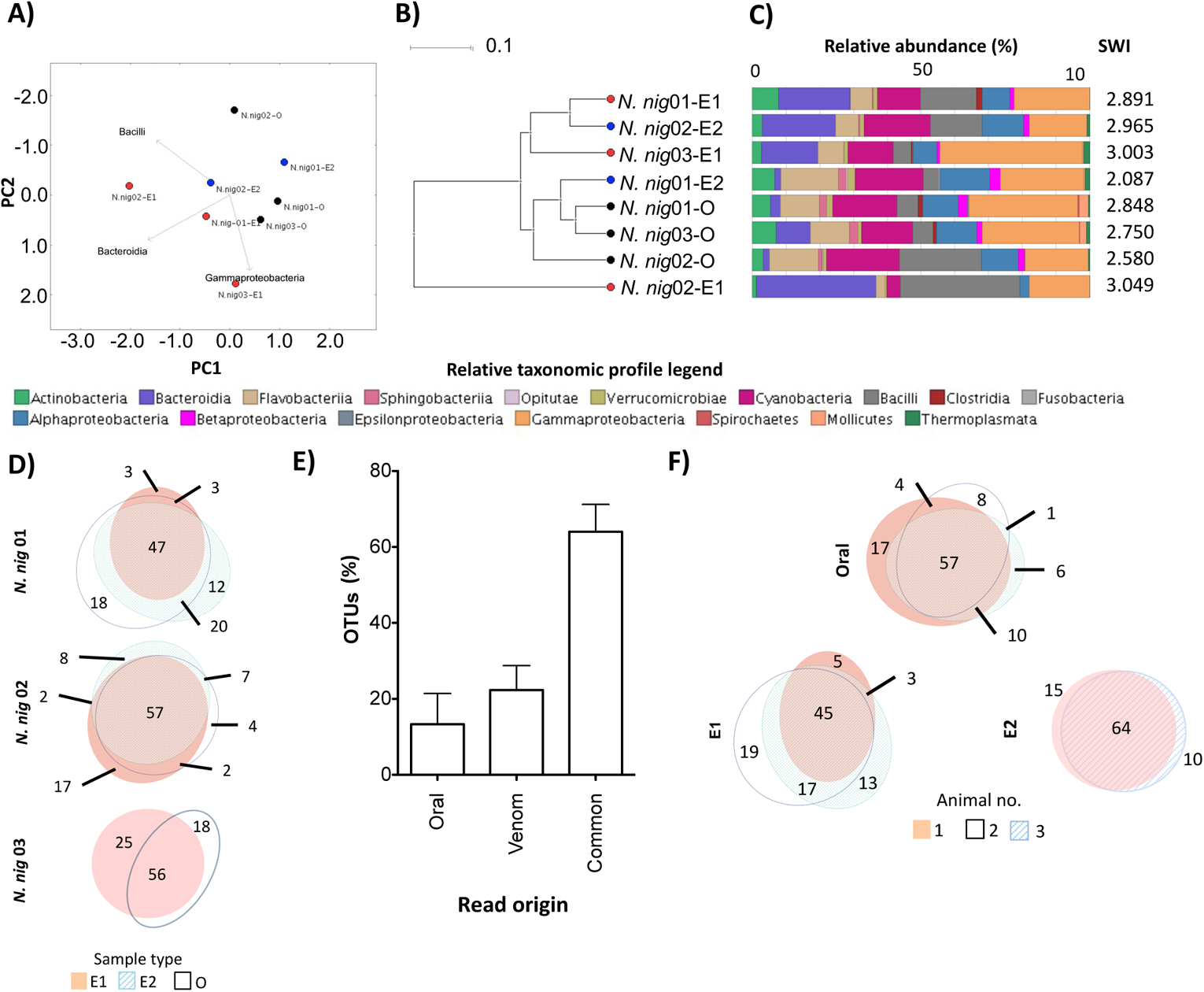
The intra- and inter-individual relationship of venom and oral microbiomes in *N. nigricollis*. Comparison of the oral and venom microbiomes in three *N. nigricollis* individuals by A) PCoA, B) UPGMA tree and C) class-level taxonomic profiling following 16S rRNA phylogenetic analysis indicates separate clustering of the microbiotae in the two microenvironments. D) Within animal incidence comparisons of operational taxonomic units (OTUs) suggest E) unique taxa exist within the oral but also the venom microenvironments. F) Between animal comparisons per niche (E1, E2, Oral) indicate most OTUs are shared but some OTUs are unique to each animal for each site. Dots in (A) and (B) represent individual *N. nigricollis* (N.nig) animal data and are coloured/labelled by sample type (black: oral; red: envenomation 1 (E1); blue: envenomation 2 (E2)). Relative taxonomic diversity profiles in (C) are aligned to the UPGMA tree sample labels, with the Shannon-Weiner Index (SWI) of each sample indicated. The ‘venom’ histogram in E) represents the sum OTU fraction found in the two envenomation samples per individual (+/- standard deviation).

### The venom flora in snakes and spiders is viable

Testing microbiome viability on identification agar (**Supp. Table 1**) yielded less growth with swab samples. Where this was significant, it was not usually matched by similar growth from the corresponding venom samples, further suggesting that the venom bacteria were probably not mouth contaminants. Strikingly, substantial and consistent growth was encountered amongst *N. nigricollis* (**Fig. 3A)** and *P. regalis* (**Table S1**) samples on blood agar. Unexpectedly, neither the wild (air dried) nor the commercial (lyophilised) venom samples yielded any growth, although colonies were obtained in blood agar from the captive *B. arietans*, underscoring the impact of venom handling on microbiome viability.

**Figure 3:**
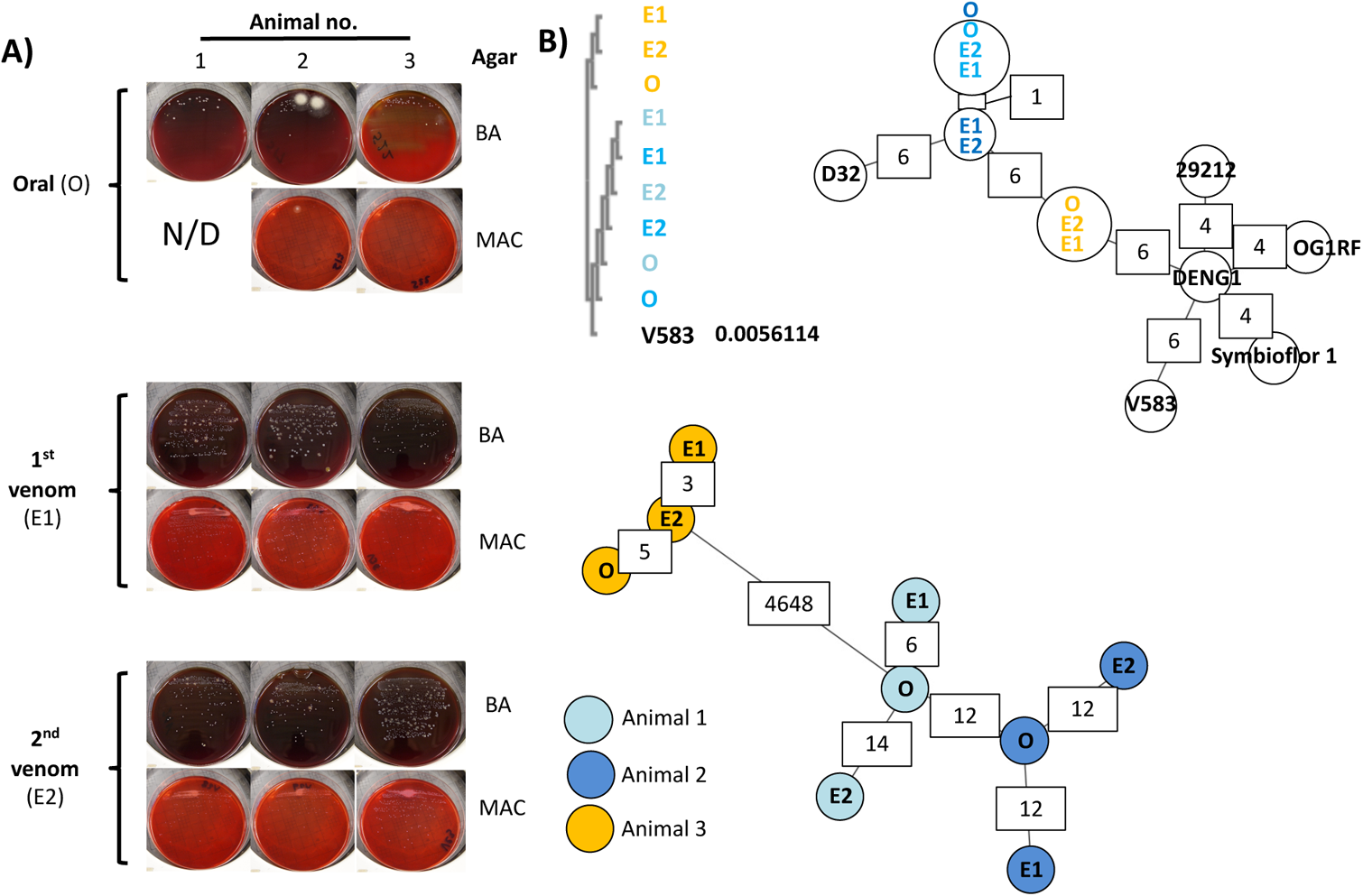
Whole genome sequencing identifies viable bacteria in *N. nigricollis* venom as two, animal-specific, *E. faecalis* strains. (A) White punctate colonies were recovered in blood agar (BA) and MacConkey agar (MAC) blinded cultures of individual oral swab (O) and two consecutive envenomation samples (E1 and E2) obtained from three captivity *N. nigricollis* snakes. N/D: none detected. (B) Blinded multiple sequence alignment (ClustalO followed by ClustalW phylogeny) of homologous sequences across the *de novo* assembled genomes against the *E. faecalis* V583 *KatA* gene (distance to V583 *KatA* indicated in V583 track) suggests two sequence groups reflecting the history and housing of the host animals. (C) Blinded MST construction based on the MLST of the *N.* nigricollis-derived isolates against nine *E. faecalis* reference genomes again separate samples into two distinct clusters that reflect the history and housing of the host animals. Partially available allele data are included in this analysis and allelic difference instances between nearest neighbours are annotated in white boxes. (D) Blinded complete genome MLST against a custom schema generated using three closely related *E. faecalis* reference genomes cluster these isolates by animal of origin (animals 1 (light blue), animal 2 (dark blue) and animal 3 (orange)). The host animal colour scheme depicted in D) is also used in B) and C).

Well-established clinical microbial biochemistry tests identified the multiple, punctate white colonies from *N. nigricollis* almost universally as *Staphylococcus spp.*, albeit with assay confidence intervals (CI) below 50% (**Table S2**). In contrast, *Stenotrophomonas maltophila* (80.4% CI) was present in five out of six *P. regalis* (all animals positive) and two *Lasiodora parahybana* (salmon pink tarantula) venom samples, but not on any fang swabs. Perplexed by the *N. nigricollis* results we sequenced these isolates on the Ion Torrent PGM.

### Viable bacteria in *N. nigricollis* venom are two new *E. faecalis* sequence types

Resequencing against putative reference genomes (**Table S2**) demonstrated less than 6% base alignment across all isolates. Instead, BLASTn analysis of the largest contig per isolate after *de novo* assembly identified *E. faecalis* V583 as the closest likely relative, resulting in >80% base alignment, at an average coverage of 51.2x. This was puzzling given the catalase positive isolate biochemistry vs. the generally accepted catalase negative nature of *E. faecalis* (19). However, the *E. faecalis* V583 *katA* gene, previously reported to encode a haem-dependent cytoplasmic catalase (20), was confirmed by BLASTn amongst all isolates at 99% identity, explaining the biochemical misclassification. Blinded multiple sequence alignment (MSA) further revealed two *katA* alleles: one shared between isolates from animals 1 and 2 (allele 1) vs. another found in animal 3 isolates (allele 2; **Fig. 3B**) varying by less than 20 single nucleotide polymorphisms to the V583 allele (**Fig. S5A**). Interestingly, these alleles grouped isolates according to the origin and joint housing histories of animals 1 and 2, vs animal 3.

To explore isolate relationships further we generated minimum spanning trees (MST; **Fig. 3C**) by multi-locus sequence typing (MLST; **Table 2**), including at core genome level (cgMLST; **Fig. 3D** and **Fig. S5B-D**) (21, 22). Comparisons to five complete reference genomes of the closely related *E. faecium* succeeded only for the *gyd* (alelles 16, 19) and *adk* (allele 18) loci. In contrast, MLST succeeded for all *E. faecalis* loci (**Table 2**), grouping isolates in line with the *katA* allele observations (**Fig. 3B**), and identifying two novel sequence types featuring two new alleles for *pstS* and *yqiL* (**Fig. S6**) as confirmed by Sanger sequencing. MLST also indicated closer relationships to the *E. faecalis* strains OG1RF, D32, and DENG1, with 87.5% +/- 1.7 of OG1RF cgMLST targets accepted for distance calculations vs D32 (78.8% +/- 2.1) and DENG1 (77.4% +/- 1.5). Pairwise comparisons of the resulting custom cgMLST schema including 5041 loci found across all the *N. nigricollis-*derived isolates also grouped isolates by their host animal (**Fig. 3C**) in line with the *katA* and MLST locus observations. Collectively, these results suggest independent acquisition from the environment of two separate and novel *E. faecalis* strains across these three animals.

**Table 2:** Novel sequence types of *E. faecalis* recovered from fangs and venoms of *N. nigricollis*.

| Animal no. | Sample |  | Locus |  |  |  |  |  |  |
| --- | --- | --- | --- | --- | --- | --- | --- | --- | --- |
|  | Isolate origin | Blinding code | *gdh* | *gyd* | *pstS* | *gki* | *aroE* | *xpt* | *yqil* |
| 1 | O | S22 | 22 | 6 | 31 | 13 | 11 | 35 | 8 |
|  | E1 | V36 | 22 | 6 | 31 | 13 | 11 | 35 | 8 |
|  | E2 | V29 | 22 | 6 | 31 | 13 | 11 | 35 | 8 |
| 2 | O | S17 | 22 | 6 | 31 | 13 | 11 | 35 | 8 |
|  | E1 | V31 | 22 | 6 | 31 | 13 | 11 | 35 | 8* |
|  | E2 | V28 | 22 | 6 | 31 | 13 | 11 | 35 | 8** |
| 3 | O | S3 | 18 | 1 | New | 24 | 83 | 47 | New |
|  | E1 | V33 | 18 | 1 | New | 24 | 83 | 47 | New |
|  | E2 | V23 | 18 | 1 | New | 24 | 83 | 47 | New |

O: oral swab sample
E1: envenomation 1
E2: envenomation 2
\*: single base pair deletion in NGS data not validated by Sanger sequencing
\*\*: homopolymer single base extension not validated by Sanger sequencing

### Pangenomic evidence of *E. faecalis* isolate adaptation to venom

Including in cgMLST comparisons an additional 3060 loci found in some, but not all of the isolates (**Fig. S5D**) identified between 290 and 831 allelic differences occurring within each animal. Furthermore, whilst 80.9% of alleles varied between the two nearest neighbour isolates from the two strains, venom isolates from animals 1 and 2 were divergent by 7.15 – 10.3% to their oral isolate counterparts. Given the well-described plasticity of the *E. faecalis* genome (OG1RF: 2.74 Mb vs V583: 3.36 Mb), we next examined mobile genetic element divergence.

Detecting the *repA-2* gene from plasmid pTEF2 (**Table S3**) amongst all isolates in accordance to MLST-derived isolate groupings suggested only plasmid fragments were found in these genomes. However, as pTEF2 is one of three *E. faecalis* V583 plasmids associated to vancomycin resistance (23), we confirmed fragments from all three pTEF plasmids (**Fig. 4A)**, consistent to MLST profiles (**Table S4**, **Table 2**), and with some pTEF sequence elements not found in secondary envenomation isolates from animals 1 and 2. Thus, the average per base read pTEF1/pTEF2 ratios for isolates from animal 3 were 0.25 (+/- 0.01), as compared to 0.90 (+/- 0.15) and 0.81 (+/-0.05) for isolates from animal 1 and 2, respectively; similarly, the pTEF2/pTEF3 average per base read ratios were 0.68 (+/-0.02) for isolates from animal 3 vs. 1.57 (+/-0.04) for isolates from animal 1 and 2. Most importantly, however, many of the genomic elements with high (>95%) sequence identity to these plasmids were also known, highly mobile sequences common to other plasmids (e.g. the *E. faecalis* Bac41 bacteriocin locus) (24). These results therefore indicated the presence of highly mobile sequences in these isolates, either on plasmids or integrated in the bacterial chromosome, that appeared to participate in the observed genomic divergence of *E. faecalis* within each animal.

**Figure 4:**
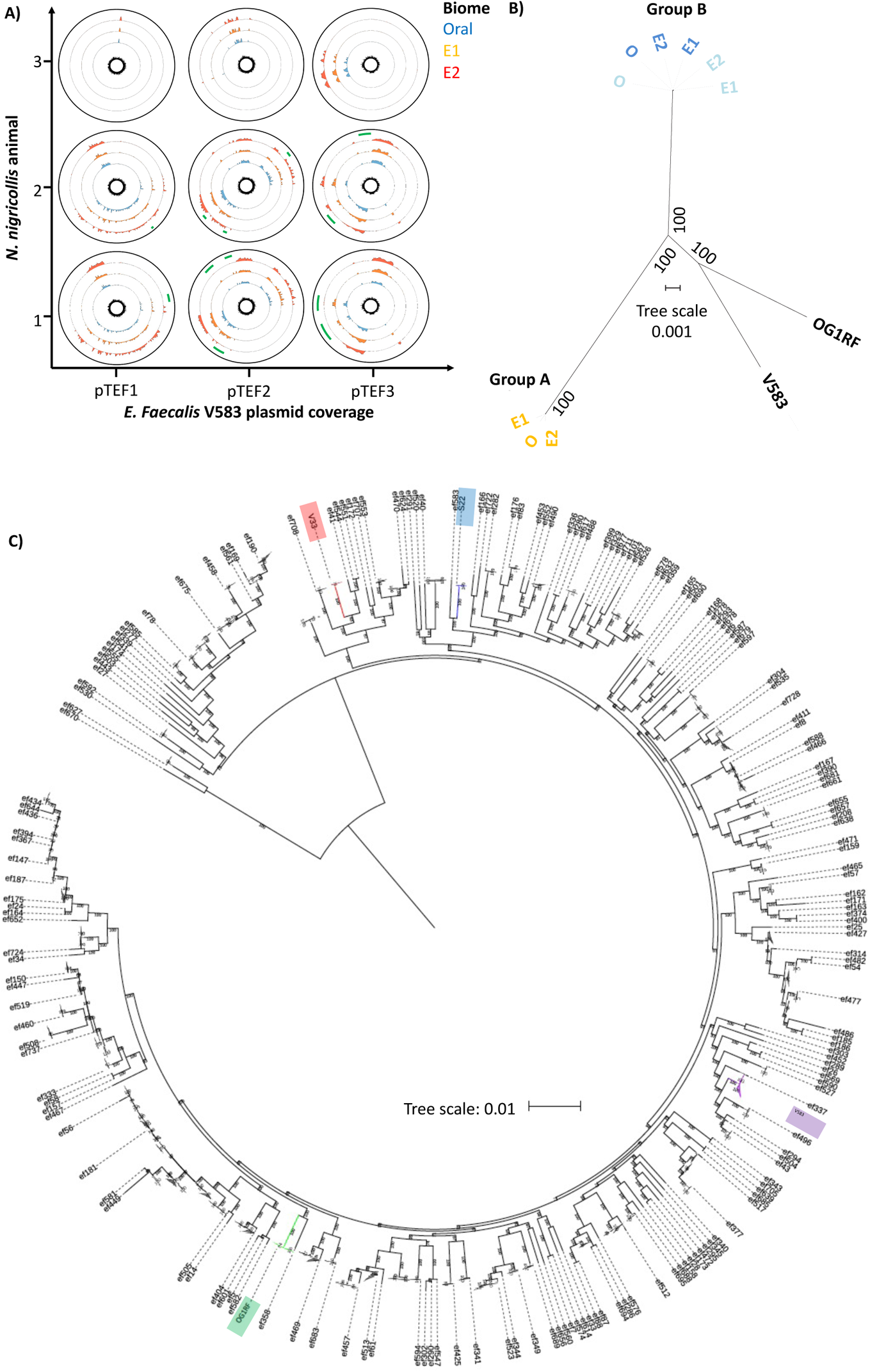
Comparative genomics of mobile and core genomic chromosomal elements of venom-tolerant *E. faecalis*. (A) Circos coverage plots of the vancomycin resistance-associated, V583 plasmids pTEF1, pTEF2 and pTEF3 in the *E. faecalis* isolates obtained from oral, envenomation 1 (E1) and envenomation 2 (E2) samples from three *N. nigricollis* individuals, reinforce the two sequence type groupings and highlight within animal variation (green arcs) indicative of sample-specific variation (lack of reads) across E2 samples in animals 1 and 2. The central plot for each plasmid and animal reflects GC content. All data is represented in 50 nt blocks. (B) Blinded maximum likelihood tree of the core genomic alignments for the 6 *N. nigricollis-*derived *E. faecalis* isolates against the V583 and OG1RF reference strains, with colour-coding refering to the origin of the isolates; light blue: animal 1; dark blue: animal 2, yellow: animal 3 (C) A maximum likelihood tree from concatenated nucleotide sequence alignment of 865 core genes (381,319 bp) from 734 genomes after removing the sites with gaps. Best-fit GTR+I+G4 substitution model was used with 100,00 ultra-fast bootstraps and SH-aLRT tests. The tree was re-rooted on the longest branch and branch lengths <0.001 were collapsed. The scale bar shows number of nucleotide substitutions per site. Branches in red, blue, purple and green show Group A, Group B, and clades containing strains V853 and OG1RF, respectively.

Closer examination of the draft genome assemblies indicated that the three *E. faecalis* isolates from animal 3 shared close genomes of ~2.9 Mb with 2,772 to 2,836 genes. The rest of the genomes varied from 3.04 to 3.24 Mb in size and encompassed 3,128 to 3,282 genes (**Table S3**). The *E. faecalis* pan-genome, including the strains OG1RF and V583, contained a total of 5,130 genes of which 1,977 belonged to the core genome. As with other analyses, the core genomic diversity separated the snake-derived strains into two major groups (**Fig. 4B**): three isolates with smaller genome sizes corresponded to animal 3 forming group A, and the remaining six isolates with larger genome sizes forming group B. However, in line with cgMLST data (**Fig. 3C**) OG1RF and V583 were quite distinct from both of these groups. Comparison of the annotated genomes indicated that 235 genes that were specific to group A and were absent from group B isolates and 321 genes specific to group B. Most interestingly, examination of the nine genomes from the snake-derived isolates identified 45 genes unique to them and absent from the OG1RF and V583 reference genomes. In contrast to Group A- and Group B-specific genes, 46.7% of these 45 genes were of known function (**Table S5**). Moreover, UniProt functional annotation indicated that 15 of these genes (35.7%) were associated to cell wall/membrane integrity, with four additional genes (9.53%) associated to pathogenic foreign protein and toxin defence.

Significantly, pathway analysis via DAVID using *B. subtilis* orthologues identified significant enrichment (8 fold, p=0.018) in the two component system pathway, specifically genes responsive to cationic antimicrobial peptides, cell wall active antimicrobials and bacitracin efflux.

To ascertain the origin of these isolates, we executed a further comparison of venom-tolerant strains with 723 additional *E. faecalis* genome sequences obtained from GenBank. As a result, the size of the pangenome increased by 5-fold (26,412 genes) with only 342 genes highly conserved among all 734 strains. A total of 865 genes were present among >99% strains. A maximum-likelihood tree from the core genome alignment separated the venom-tolerant strains into different groups with isolates from diverse sources globally including animal, environmental and human isolates (**Fig. 4C; Supplementary Fig. S7**). However, the venom-tolerant strains formed distinct subclades within large groups (**Supplementary Fig. S7**).

Genome comparison identified 42 genes that were unique to group A and were absent in other *E. faecalis* genomes (**Table S6**). However, 18 of these genes encoded hypothetical proteins.

Similarly, 97 genes were only observed in group B with 65 genes encoding hypothetical proteins, or proteins of undefined functions (**Supplementary Table S6**). Although these genes appear to be specific to the venom-tolerant groups based on the 70% identity threshold used in the genomic comparison, homologs or proteins with the same function may be present in other strains. Indeed, genes with similar functions were observed for most of the group-specific proteins with an annotated function in strains outside the group (**Table S7-8**). A group-A gene, S3_02356 encoding a colanic acid biosynthesis protein did not return any hits but a gene (ef95_02851) with the same function was also present in *E. faecalis* strain 7330112-3. Similarly, four genes (S22_00125, S22_03166, S22_03201 and S22_03202 encoding CAAX amino terminal protease self-immunity, Biotin transporter BioY, Bacterial regulatory proteins of gntR family and PRD domain protein, respectively) did not return hits based on our search criteria, but genes with the same functions were widely present in the dataset. Any potential role of these genes and their associated pathways in adaptation or tolerance to venom will require further molecular characterisation in the future.

Snake venoms often contain metalloproteinases, phospholipases A2 (PLA2), serine proteases, three-finger peptides (3FTX) and other secretory proteins including L-Amino acid oxidase (25, 26). *N. nigricollis* venom is cytotoxic due to the presence of PLA2 and 3FTX that are involved in heamolysis and methylation of haemoglobins causing severe hypoxia in humans with fatal consequences due to cardiovascular and neurological failure (27). PLA2, and 3FTXs facilitate disruption of membrane integrity in bacteria (28, 29). Two genes encoding LPL transporter (LplT) and acyltransferase-acyl carrier protein synthetase (Aas) were found to protect bacterial cell envelope from Human PLA2 in Gram-negative bacteria (30). In addition, mutations in cell wall (*dltA*) and cell membrane (*mprF*) polyanions are found to be involved in increased sensitivity to human PLA2 in *S. aureus* (31). LplT is only observed in Gram-negative bacteria but homologs of Aas, DltA, and MprF were detected among the venom isolates in this study. The *mprF* gene appears to be disrupted in strain V31 which is annotated as two smaller proteins (V31_01061 and V31_01062; **Table S9**). Sortase A has also been associated with the resistance to human PLA2 in *Streptococcus pyogenes* (32) and 3 copies of sortase family proteins was detected among all 9 venom isolates. Therefore, all 9 isolates may be able to maintain integrity of the cell membrane in presence of venom.

L-Amino acid oxidase can inhibit bacterial growth of both Gram-negative and Gram-positive bacteria by exerting oxidative stress due to the release of reactive oxygen species during oxidative deamination of l-amino acids (25). 21 genes involved in oxidative stress response are present among venom-tolerant strains, including genes encoding various antioxidative enzymes such as superoxide dismutase, catalase, and glutathione metabolism that are known to protect cells from free radicals and also contribute to the virulence of *E*. *faecalis* strains (**Table S9**; (33–35). Therefore, these *E*. *faecalis* strains appear to be well-equipped to survive the stress imposed by *N. nigricollis* venom. This tenet was further supported experimentally, wherein *E. faecalis* V583 growth was dose-dependently inhibited at a minimum concentration of 11.7 mg/ml (95% CI: 9.36-14.6 mg/ml) and non-inhibitory concentration of 2.78 ng/ml (95% CI: 2.21-3.50 mg/ml) of filter-sterilised freeze-dried *N. nigricollis* venom in brain heart infusion broth. By stark contrast, all the venom-derived strains exhibited <30% growth inhibition even at 50 mg/ml freeze-dried venom concentrations i.e. ~4x lower than the 208 mg/ml concentration of fresh *N. nigricollis* venom, resulting in ambiguous non-inhibitory concentration ranges of 25.2-44.0 mg/ml (no CI’s calculable), with the group A strain V33 exhibiting no susceptibility to the inhibitory effects of venom **(Fig. S8)**.

### Projecting primary infection clinical risk from venom-tolerant *E. faecalis* isolates

Given these viable *E. faecalis* strains could potentially infect envenomation wounds, we next examined the genomic data for known resistance determinants that might facilitate opportunistic primary infection. None of these strains had any acquired resistance genes to any antimicrobial classes (**Table S10**), although all isolates featured *lsaA*, which confers intrinsic streptogramin resistance to *E. faecalis* (36): MSA further reinforced isolate grouping based on host animals (**Fig. S9**). Since horizontally acquired genes are largely responsible for resistance to vancomycin, aminoglycosides, macrolides, and tetracycline in *Enterococcus* (37), these strains were considered to be susceptible to drugs in each of these drug classes. In addition, the absence of known resistance-associated mutations in *gyrA*, *parC*, and the 23S rRNA genes also increased the likelihood that these isolates would be susceptible to oxazolidinones and fluoroquinolones.

Thus, these data indicated that several available antimicrobials would likely be effective in treating infections caused by these strains of *E. faecalis*. However, a gene related to macrolide export (macB5) was detected in the venom microbiome strains (**Table S5**).

On the other hand, one additional concern was the potential misidentification of these isolates as *Staphylococcus* using standard diagnostic methods: this could potentially impact treatment decision-making. Although many of the same antibiotics, including vancomycin and linezolid, would be considered for treatment of both staphylococci and enterococci, there are some potential differences. For instance, oxacillin is often employed as a first-line agent to treat *Staphylococcus* (38). This drug is not effective against enterococci, as the use of penicillins for *E. faecalis* infections would typically involve ampicillin, usually in combination with an aminoglycoside (39). In addition, cephalosporins such as cefotaxime are considered second-line therapies for coagulase-negative staphylococci such as *S. epidermidis* (40). However, enterococci are intrinsically resistant to this class of drugs, and their prevalence in the gut actually tends to increase in response to cephalosporin therapy (41). Thus, while the *E. faecalis* strains in this study did not have any known acquired resistance determinants, if they were to cause infections, ensuring their proper identification would be critical to issuing correct treatment and achieving positive clinical outcomes.

## Discussion

In contrast to the generally held view that venoms are both antimicrobial (1–4) and sterile (6, 7, 42), despite contrasting reports since the 1940’s (43), we show that microorganisms are common and viable in the venoms of both vertebrates and invertebrates. As with previous work (11–13), our data support the precept that prey faeces might seed the oral and the venom microbiome.

However, significant adaptation takes place in these bacteria to persist in venom, occurring in parallel within each animal probably through horizontal gene transfer, to contribute genes unique to isolates from venom samples. An unusually large fraction of these genes (45.2%) is involved in maintaining bacterial membrane integrity or toxin defence. Notably, disruption of membrane integrity appears to be the most common mechanism of action for known, venom-derived antimicrobial peptides and enzymes (4, 5) of long-standing (2) and significant biotherapeutic interest, including MDR (1) and nociception (16). It is unclear at present to which extent this form of parallel convergent evolution extends beyond *Enterococcus* or other antimicrobial resistance mechanisms, such as antibiotic resistance genes against last resort antibiotics found on mobile genetic elements (44, 45) across multiple continents. This work therefore adds to the body of evidence (46) supporting further scrutiny of host-microbe interactions in the venomous microenvironment in understanding microbial adaptation mechanisms to extreme environmental challenges.

From a clinical perspective, identification of *E. faecalis* as the most prevalent culturable microbe across our European *N. nigricollis* venom samples strikingly reflects three independent clinical reports across Africa and Asia that this non-sporulating microbe is the most common Gram positive infection cultured from infected envenomation wounds (8–10). Likewise, *E. faecalis* were found to be the most common aerobic Gram-positive isolates in *N. naja* oral swabs (n=6), among a number of multidrug resistant Gram-negative and Gram-positive bacteria, including human pathogens known to cause fatal infections such as *Salmonella* enterica serovars Typhi and Paratyphi A (49). Further epidemiological data from countries with high envenomation incidence (8–10, 47, 48) challenge the consensus view in developed nations that venom is sterile, that opportunistic primary infection upon envenomation is uncommon, and that venom wound infection is a consequence of poor hygiene or poor debridement practice (6, 7). The incidence of post-envenomation infections likely to be caused by venom-tolerant bacteria is also not restricted to snakes alone. In fact, post-envenomation cellulitis and dermatitis, presumed bacterial in nature, was observed in 25% of a 16-case series of *Steatoda nobilis* (false widow spider) envenomations in the UK and Ireland: one of these required intravenous penicillin and flucloxacilline treatment after hospital admission (50). *S. nobilis* chelicerae, were previously found to harbour 11 bacterial taxa and 22 separate bacterial species, including class 2 pathogens; 3 of these 22 species showing multi-drug resistance (51). Although explicit genomic evidence connecting venom microbes to envenomation infection remains elusive, in an experimental rabbit model of dermonecrosis (48) caused by *Loxosceles intermedia* (recluse spider) venom, *Clostridium perfringens* recovered from the spider fang and venom enhanced disease symptoms.

*Stenotrophomonas-*like bacteria were also found to dominate cone snail venom microbiomes (52), indicating that microbial venom adaptation may extend well beyond snakes, spiders, scorpions, and snails. The cone snail study also reported comparable microbiomes in samples collected across the Pacific basin as well as Atlantic specimens. Furthermore, building upon the few instances of polymicrobial infection reported clinically (8–10), the reports on *L. intermedia* (48)*, Conus* (52), and herein suggest that diverse microbes effectively co-colonise venom glands in host species-specific manner, and thus envenomation wounds. Taken together, these studies support a review of the current standards of care for envenomation wound management (42) beyond simply managing the severe tissue damage and necrosis that might be caused by venomous bites, to include clinical microbiology on envenomation wounds upon presentation.

This would be particularly relevant to individuals immunocompromised through disease or malnutrition, e.g. in developing nations where envenomation incidence rates are high, or to children on a venom/colony forming unit dose per body weight basis.

Yet common microbial diagnostic methods mistook *E. faecalis* for *Staphylococcus*, which could lead to unfavourable clinical decision making. It is unclear at present how frequent such misidentification events might be, contributing to MDR through inappropriate antibiotic use. At least one retrospective study reported higher incidence of *Staphylococcus spp.* in envenomation wounds (12), and Blaylock’s seminal snake oral flora studies also reported *Proteus* and *Staphylococcus* (14): both relied on the same methods we used in this study that were found to misidentify the pathogen. Our results therefore further support use of PCR/sequencing methods as they become more relevant to resource limited settings (53), and suited to the point of need (54), in line with World Health Organisation ASSURED criteria. Understanding the sensitivity of these methods will be crucial in their reliable implementation in envenomation care. It is therefore noteworthy that despite the limited biomass levels in these samples, species-level OTU analysis on MG-RAST (55) correctly identified *E. faecalis* as one of the principle aerobic isolates in *N. nigricollis* venom. Thus, a simple phylogenetic or metagenomic approach, combined with local herpetogeography knowledge, could quickly and accurately inform clinical action regarding antivenom administration. This is because, unlike incidents involving venomous pets, envenomating animal capture is both rare and contraindicated to minimise further envenomation injuries (9). Culprit animals are also commonly misidentified through inadequate description (9), and antibody-based venom identification kits have so far proven unreliable (47), generally complicating antivenom selection.

To conclude, we evidence that vertebrate and invertebrate animal venoms host diverse, viable microbiomes, with isolates genetically adapted to venom antimicrobials of medical interest against MDR. These results challenge perceptions on the sterility of venom and absence of primary infection risk upon envenomation, pointing to modern nucleic acid technologies to better inform envenomation care and antibiotic use.

## Materials and Methods

### Animals and sampling

All samples analyzed in this study were provided by Venomtech Ltd., with the exception of freeze-dried *B. arietans* venom (Latoxan, Portes les Valence, France) and field collected samples collected in South Africa. Briefly, captive animals were housed in 2 m by 1m wooden, glass fronted vivaria with large hide, thermal gradient and water *ad-libitum*. All procedures for venom collection and swabbing were approved as unregulated under the Animals (Scientific Procedures) Act 1976. Venom was collected by standard techniques; briefly snakes were restrained behind the head and presented to a collection vessel. Snakes freely bit into the vessel until envenomation was observed. Each snake was presented to two sterile collection vessels in succession, one for the first envenomation with potential fang plug, and the other for the second flow (labelled E1 and E2, respectively). While the snake was positioned over the second vessel, the oral cavity was swabbed with a sterile swab with individual collection tubes (invasive sterile swab with transport media, DeltaLab, VWR, Lutterworth, UK). The venom collection vessels were clear, sterile 125 ml polypropylene containers (ThermoFisher Scientific Ltd., Paisley, UK) covered by 2 x 9 cm^2^ pieces of parafilm stretched to fit (ThermoFisher Scientific Ltd.). The collection vessel was secured to a bench during collection. Samples collected in the field were from wild puff adders sampled as part of a previous phylogeographic study (56). Venom samples were collected using a similar method to that described for captive animals, except that the entire venom sample was collected in a single collection vessel. Samples were lyophilised by storing <100 μl venom aliquots in a vacuum-sealed container that was half-filled with silica gel. Following drying, venom samples were stored in a refrigerator at 5°C.

*Lasiodora parahybana* and *Poecilotheria regalis* were housed in 5 and 8 litre polypropylene boxes (Really Useful Products Ltd, Normanton, UK), respectively, with moist vermiculite (Peregrine Livefoods Ltd., Ongar, UK), plastic hide and 5 cm water bowl, as previously described.^58^ Arachnids were anaesthetised with a rising concentration of carbon dioxide, the fangs were swabbed with a sterile swab which was then placed in an individual 1 ml sterile, DNA free, polypropylene collection tube (FluidX Ltd, Nether Alderley, UK), and venom was subsequently collected from arachnids by electrical stimulation. All samples were stored at −80°C. The same transport swabs (VWR) as those used for snakes were also used for invertebrate oral / aculear swabbing. Samples were stored at 4°C and cultured within 24 hours of collection.

### Microbial culture

Aerobic microbial viability was determined by plating swabs or aliquoting 10 μl volumes of venom samples onto oxalated whole horse blood agar, MacConkey agar, or mannitol salt agar (ThermoFisher Scientific) plates and incubating at 30°C for 72 hours. Biochemical isolate identification was undertaken using API^®^ strips (20E, 20NE and Staph) interpreted via the APIWEB interface (BioMerieux, Basingstoke, UK). All isolates were stored on beads at −80°C at the University of Westminster microbial isolate library. *Naja nigricollis* sub-culture was performed by restoring cryogenically stored bacteria on lysogeny broth agar (ThermoFisher Scientific) grown for 48 hours at 30°C, and single colony overnight culture in lysogeny broth (ThermoFisher Scientific) using aerated culture (300 rpm). Minimum inhibitory concentrations and non-inhibitory concentrations were determined in 96 well format assays according to Lambert *et al.* in brain heart infusion media by measuring absorbance at OD_600_ on a Tecan Spark Cyto 96 plate reader (Tecan, Männedorf, Switzerland). All bacterial agar and broth materials were purchased from Formedium Ltd. (Norfolk, UK).

### DNA extraction

Neat venom samples or samples diluted in 18 megaohm water previously confirmed as bacterial DNA free by 16S PCR were subjected to DNA extraction using TRIzol^TM^, PureLink Genomic DNA kits or MagMAX Cell-Free DNA kits (ThermoFisher Scientific) according to the manufacturer’s instructions. For combined extraction of Gram positive and Gram negative bacteria from liquid samples, diluted samples were split in equal volumes and processed according to the manufacturer’s Gram wall specific lysis protocols, with lysates combined prior to DNA binding onto columns by simple admixture. DNA content was then analyzed by Nanodrop (ThermoFisher Scientific) spectrophotometry and purified material was stored at −80°C until further analysis.

### 16S phylogenetic library preparation and sequencing

For short amplicon library preparation, the hypervariable V3 region of the 16S rDNA gene was amplified from 20 ng of DNA using the primers 5′-CCTACGGGAGGCAGCAG-3′ and 5′-ATTACCGCGGCTGCTGG-3′ (Integrated DNA Technologies BVBA, Leuven, Belgium),(18) 1U Platinum^®^ PCR SuperMix High Fidelity (ThermoFisher Scientific) and 10 μM of primer-mix. The reaction mixes were incubated at 94 °C for 5 min followed by 30 cycles of 30 seconds at 94 °C, 30 seconds at 55 °C and 1 min at 72 °C and then final elongation at 72 °C for 10 min using a Techne Prime Thermal cycler (ColePalmer, Staffordshire, UK). PCR products (193 bp) were confirmed by 2% w/v agarose gel electrophoresis in TAE buffer (ThermoFisher Scientific).

NGS library preparation was carried out using the Ion Plus Fragment Library Kit according to the manufacturer’s instructions (Rev. 3, ThermoFisher Scientific), except that reactions were reduced to 1/5^th^ volumes. Pooled libraries were diluted to ~26 pM for templating on the Ion OneTouch 2 system (ThermoFisher Scientific) using the Ion PGM Template OT2 200 v2 kit according to the manufacturer’s instructions (Rev. B, ThermoFisher Scientific). Templated samples were sequenced on the Ion Torrent Personal Genome Machine (PGM; ThermoFisher Scientific) system on a single 318 Ion Chip (ThermoFisher Scientific) using the Ion PGM^TM^ 200 Sequencing kit according to the manufacturer’s instructions (Rev G., ThermoFisher Scientific).

### Whole genome sequencing

DNA extracted from cultured isolates was mechanically sheared using the Covaris S220 Focused-ultrasonicator (Covaris, Brighton, UK). NGS libraries were generated using the NEBNext Fast DNA Library Prep Set for Ion Torrent (New England Biolabs, Hitchin, UK). Pooled samples were size selected with the LabChip XT (LabChip XT DNA 300 Assay Kit; PerkinElmer, Seer Green, UK) and diluted to 26 pM for templating with the Ion OneTouch 2 system using the Ion PGM Template OT2 200 kit. Templated samples were sequenced on the Ion PGM using the Ion PGM™ Sequencing 200 v2 reagent Kit (ThermoFisher Scientific) and Ion 318™ v2 Ion Chip (ThermoFisher Scientific).

### Bioinformatic analyses

Raw Ion Torrent sequencing data reads were quality controlled and demultiplexed using the standard Ion Server v. 4.0 pipeline (ThermoFisher Scientific). Referenced and *de novo* assemblies were carried out using TMAP v.4.0 and the SPAdes plugin in the Ion Server. Phylogenetic data analyses were carried out after independent data deposition and curation on the MG-RAST v.3.0 pipeline (55) (project IDs MGP5177 and MGP5617) which uses a BLAST approach and the EBI-METAGENOMICS v.1 (project ID ERP004004) pipeline (57) which uses a Hidden Markov Model approach. Raw 16S sequencing reads were deposited in the European Nucleotide Archive (PRJEB4693). Quality control for both resources included length and quality filtering followed by a dereplication step where sequences with identical 50 nucleotides in 5’ positions were clustered together. MG-RAST taxonomy annotation involved RNA identification using VSearch, and assignments using a custom database generated by 90% identity clustering of SILVA, GreenGenes and RDP prokaryotic databases. EBI-METAGENOMICS identified rRNA using Hidden Markov Models present in the RDP databases and assigned taxonomy using Qiime and the GreenGenes prokaryotic database.

For post-processing analyses, the EBI-curated dataset was analyzed using MEGAN v.5.5.3 (58). Classical multi-locus sequence typing (http://efaecalis.mlst.net/) and cgMLST (21, 22) were carried out using Ridom SeqSphere+ v.4.0 running on a 2 core, 10 GB RAM, 500 GB hard disk Biolinux v.8.0 installation on a VirtualBox virtual machine instance on a 16GB RAM, 1TB hard disk Apple iMac. Extended cgMLST analysis to include partially detected loci, excluded loci annotated as ‘failed’ due to sequencing error suggesting genuine *E. faecalis* genomic divergence occurring within each animal. Plasmid detection was carried out using the PlasmidFinder v.1.3 server (59), followed by NCBI BLASTn analysis to detect shorter fragments, e.g. the same 398 nt fragment of *repA-2* in animal 3 isolates (<40% of the full-length gene) at 90.1% identity to the plasmid-borne reference sequence. Single gene comparisons and multiple sequence analyses were carried out using TCoffe and MView on the EMBL-EBI server, with base conservation visualized by BoxShade v.3.3.1 on mobyle.pasteur.fr. Genome-level plasmid coverage analyses were carried out by NCBI BLASTn and comparisons were visualized using Circos v.0.69-4.

The sequencing reads were assembled using SPAdes v.3.9.0 (60), and the draft assemblies were annotated using Prokka (61) before NCBI deposition (BioProject No. PRJNA415175). The genome sequences of *E. faecalis* strains V583, OG1RF (Accession numbers NC_004668.1 and NC_017316.1, respectively) and 723 other *E. faecalis* strains were obtained from GenBank and were re-annotated using Prokka to have an equivalence of annotation for comparative analyses.

The genomes were compared using the program Roary with a protein similarity threshold of 70% (62, 63). A maximum-likelihood tree was constructed from the core genomic alignment using IQ-Tree (64) with 100,000 ultra-fast bootstraps and 100,000 SH-aLRT tests. The tree was visualized using Interactive Tree Of Life (iTOL) (65).

To identify acquired resistance genes, nucleotide BLAST analysis was performed on the ResFinder (66) and NCBI (https://www.ncbi.nlm.nih.gov/pathogens/) resistance gene databases using cutoffs of 50% length and 85% identity to known resistance determinants. Additional BLAST analysis was performed to identify single nucleotide polymorphisms in the quinolone resistance determining region (QRDR) of *gyrA* and *parC* (67). Additional mutational analysis was performed on region V of the 23S rRNA-encoding genes (68).

BLASTP was performed in Ensembl Bacteria (release 38), against the *E. faecalis* V583 and *E. faecalis* (GCA_000763645) to obtain further geneID’s from significant matches. *Bacillus subtilis* orthologue gene ID’s were collated as this species is the closest relative to *E. faecalis* (VetBact.org) with the most comprehensive genome annotation required for gene onotolgy and KEGG pathway analysis. From the 42 genes unique to venom isolates, useable *B. subtilis* GeneID’s were obtained for 20, of which 18 of these successfully converted to ENTREZ Gene ID’s using the functional annotation tool (DAVID Bioinformatics resource 6.8)(69, 70), selecting *B. subtilis* as the background species.

## Acknowledgments

We would like to thank: Drs. Pamela Greenwell and Caroline Smith for their invaluable input on non-standard DNA extraction methodology options suited to unusual samples; Dr. Patrick Kimmit for his input on microbial characterisation; Mr Peter Gibbens for housing and venom collection from captive *N. nigricollis* and *B. areitans*.

## Author contributions

a MMGL and TDL sampled, and CT and ST prepared the library of captive animal venoms. WW and AB collected and prepared the wild snake samples. EE, JT, PG and SAM optimized and performed the DNA extractions and 16S PCR. JT and EE performed the preliminary and main study library preps and next generation sequencing experiments, respectively. AD, HD, PK, LS, and SAM performed the phylogenetic data quality control, curation and analysis. KFR and SAM performed the microbial cultures and biochemical characterization. MKV and LU grew the *E. faecalis* isolates and performed the whole genome sequencing. MKV, KW and SAM performed the *E. faecalis* isolate genomic characterization and MLST+ analysis. GT performed *E. faecalis* resistome analysis. VS performed the *E. faecalis* isolate pangenome data reduction and ST identified the venom resistance gene ontology subset. SAM conceived the study and designed experiments together with ST. All authors contributed equally to the overall interpretation of the dataset and manuscript preparation;

## Data and materials availability

Phylogenetic data are deposited on MG-RAST (project IDs MGP5177 and MGP5617) and the EBI-METAGENOMICS servers (project ID ERP004004). Raw 16S sequencing reads were deposited in the European Nucleotide Archive (PRJEB4693). Annotated draft *E. faecalis* genome assemblies are deposited on NCBI (BioProject No. PRJNA415175).

## Supplementary Materials

Figures S1-S7

Tables S1-S8

References (41–53)

**Figure S1:**
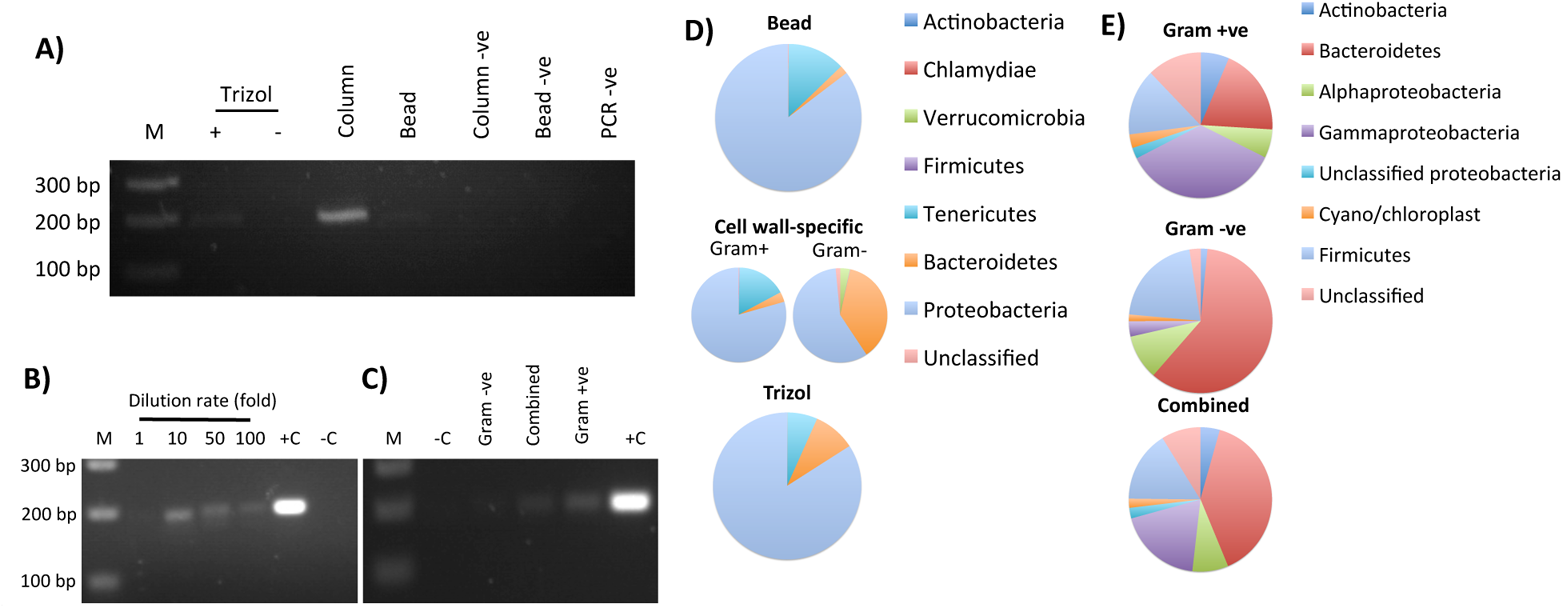
16S Ribosomal RNA gene PCR output and phylogenetic differences on account of venom collection and extraction methods. The choice of extraction method (phenol-chloroform-based: Trizol; column based; magnetic bead based) impacts significantly on the recovery and amplification of bacterial DNA from lyophilized *B. atrox* venom (A). This bacterial DNA is not an artefact of lyophilisation process contamination as detection is maintained in aseptically collected, flash-frozen *Bitis arietans* venom, nor is it an artefact of diluent contamination by 18 MΩ water confirmed 16S free by PCR; however, >10x dilution of venom is necessary for PCR to progress (B). The yield of bacterial DNA is a function of upstream cell lysis methods selectivity for Gram +ve or Gram –ve cell walls (C). The cell lysis and extraction methodology also directly impact upon microbial diversity profiles as determined by 16S rRNA phylogenetics for either lyophilised (D) or aseptically collected, flash-frozen venoms (E),, with combined use of cell wall-specific extraction methods yielding more balanced profiles.. +C: positive control; −C: negative control; −ve: method specific negative controls.

**Figure S2:**
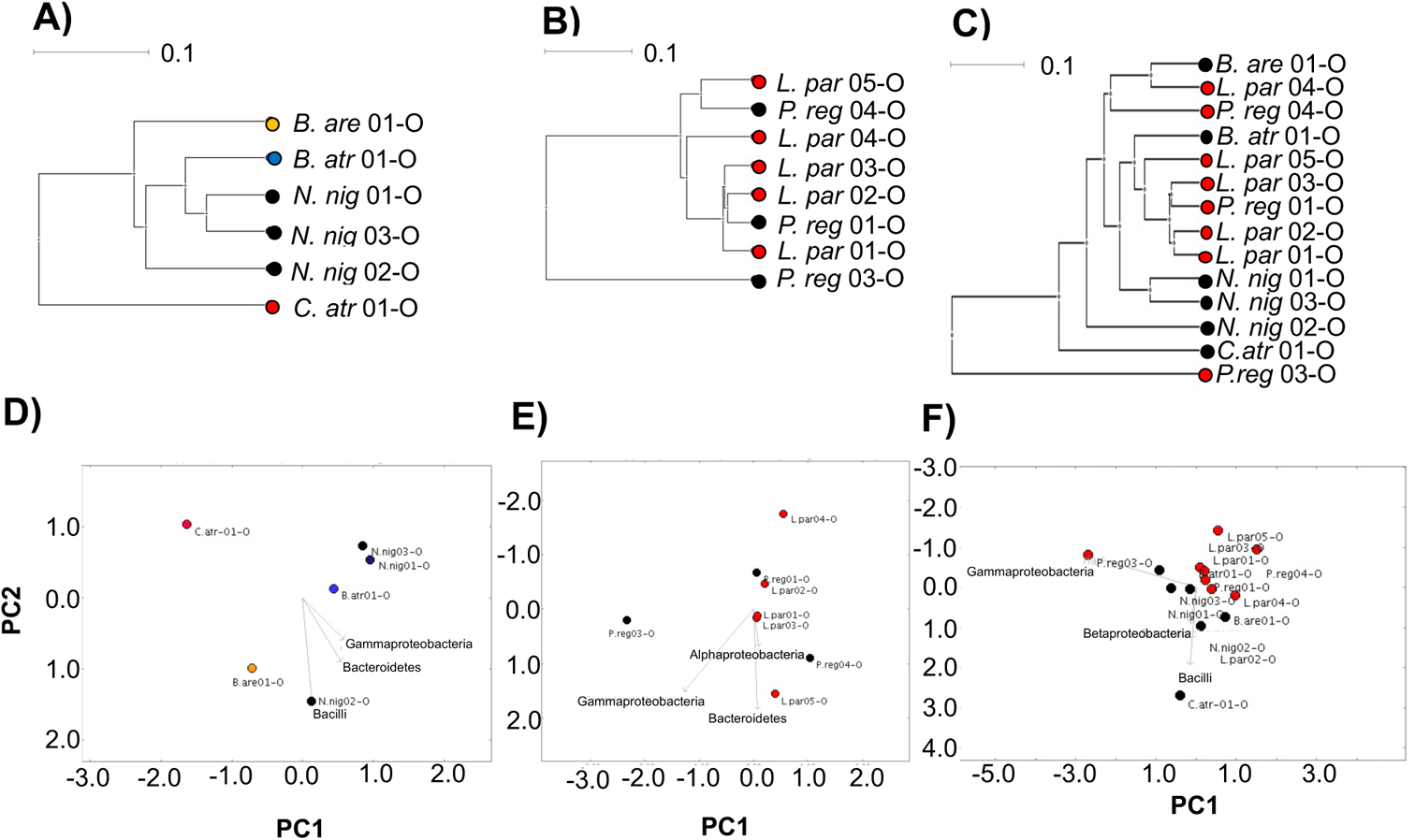
Comparison of the oral microbiomes of snakes and spiders suggests their oral microbiota is not host-species specific. UPGMA tree (A - C) and PCoA (D – F) analysis (Bray-Curtis indeces) of the oral microbiome diversity of snakes (A, D; individual species identified by independently coloured dots), spiders (B, E; *L. parahybana*: red dots, *P. regalis*: black dots), or vertebrate vs invertebrate animals (C, F; black vs red dots), as determined by 16S rRNA phylogenetic analysis at class level indicate no host species-specific relationships. Dots represent single captivity individuals, labelled with short species name, enumerated for individual number and identified for the oral/fang (O) nature of the sample.

**Figure S3:**
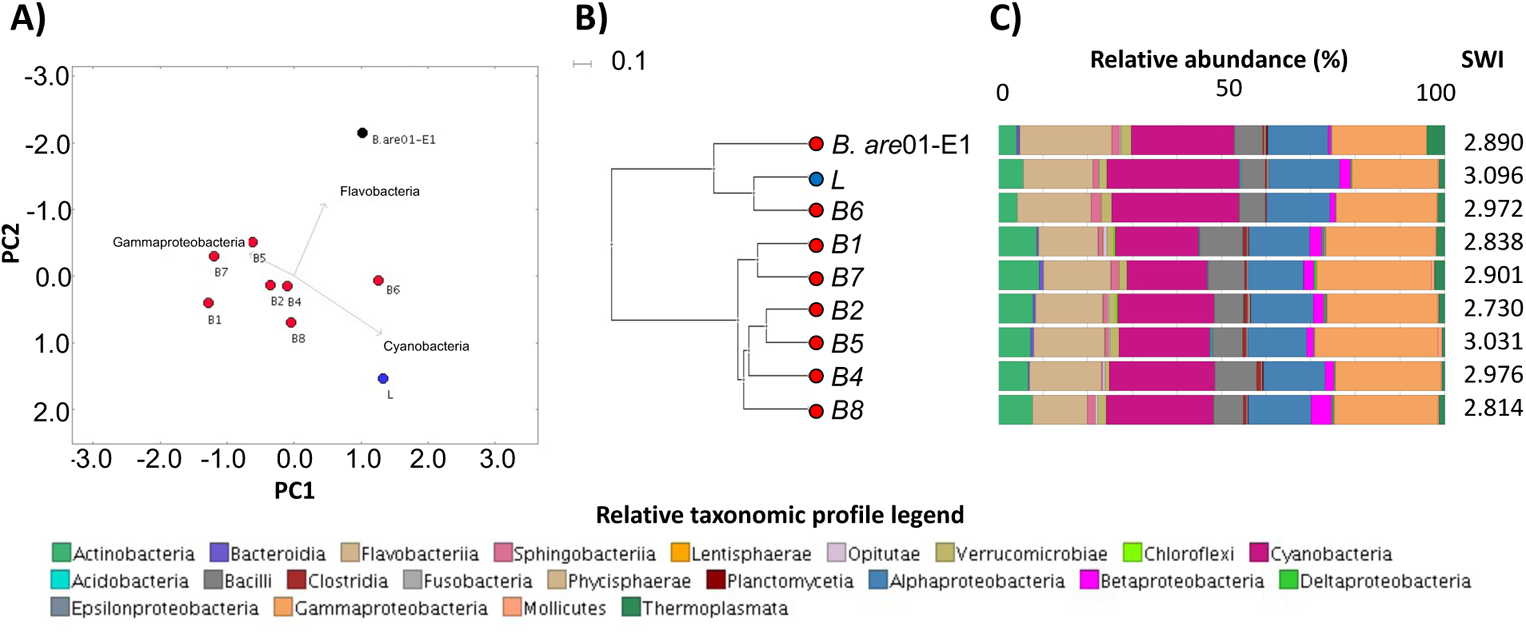
The origin of a *B. arietans* snake does not appear to influence the microbiome profile in the venom of each animal. *B. arietans* venom microbiome profiles do not present substantial differences on account of host geographical origin as determined by A) PCoA, B) UPGMA tree and C) class-level taxonomic profiling following 16S rRNA phylogenetic analysis. Dots in (A) and (B) represent individual animal data, are coloured and labelled by animal origin and number (red B1-8: wild; blue L: lyophilised captivity; black B. are01-E1: flash-frozen captivity). Relative taxonomic diversity profiles in (C) are aligned to the UPGMA tree sample labels, with the Shannon-Weiner Index (SWI) of each sample indicated.

**Figure S4:**
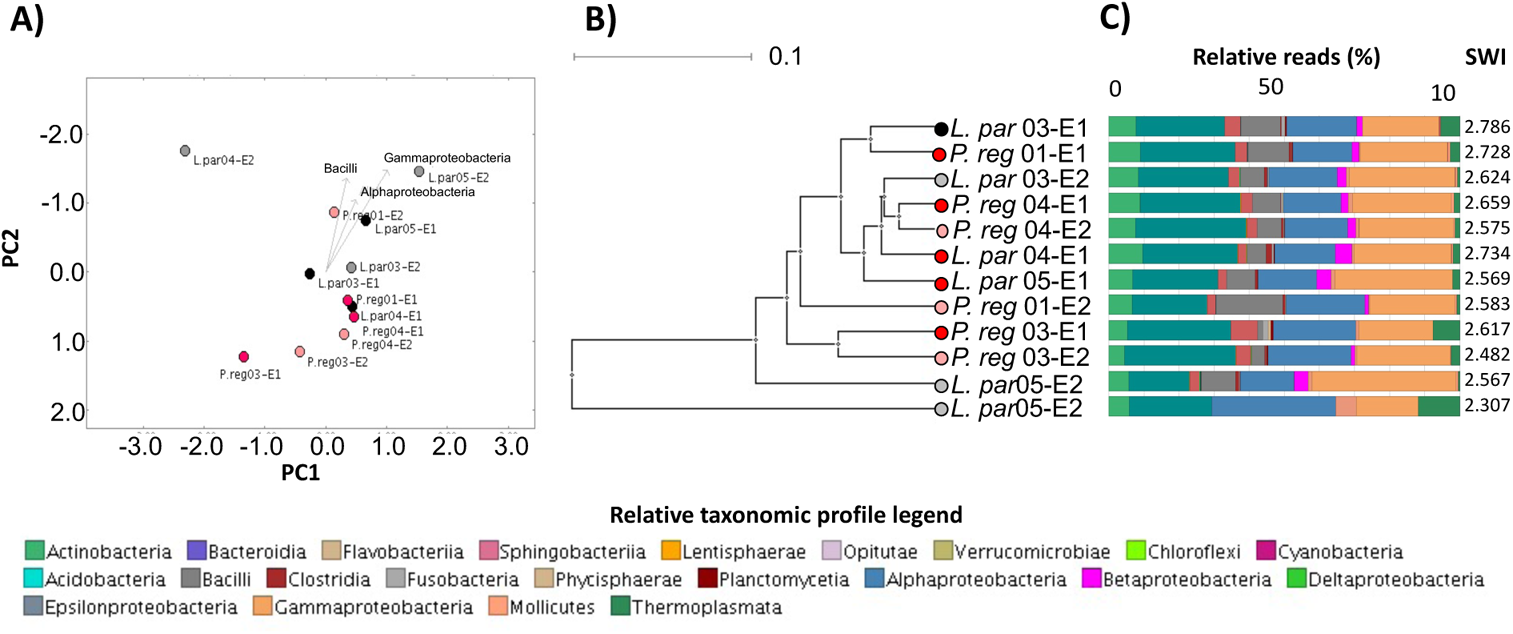
Spider venom microbiome profiles suggest closer relationships between consecutive envenomation samples within *P. regalis* individuals. Spider venom microbiomes were compared by A) PCoA, B) UPGMA tree and C) class-level taxonomic profiling following 16S rRNA phylogenetic analysis. Dots in (A) and (B) represent individual animal data, are colored/labelled by species and envenomation number (black and grey: *L. parahybana* envenomation 1 (E1) and 2 (E2) respectively; red and pink: *P. regalis* E1 and E2 respectively). Relative taxonomic diversity profiles in (C) are aligned to the UPGMA tree sample labels, with the Shannon-Weiner Index (SWI) of each sample indicated.

**Fig. S5:**
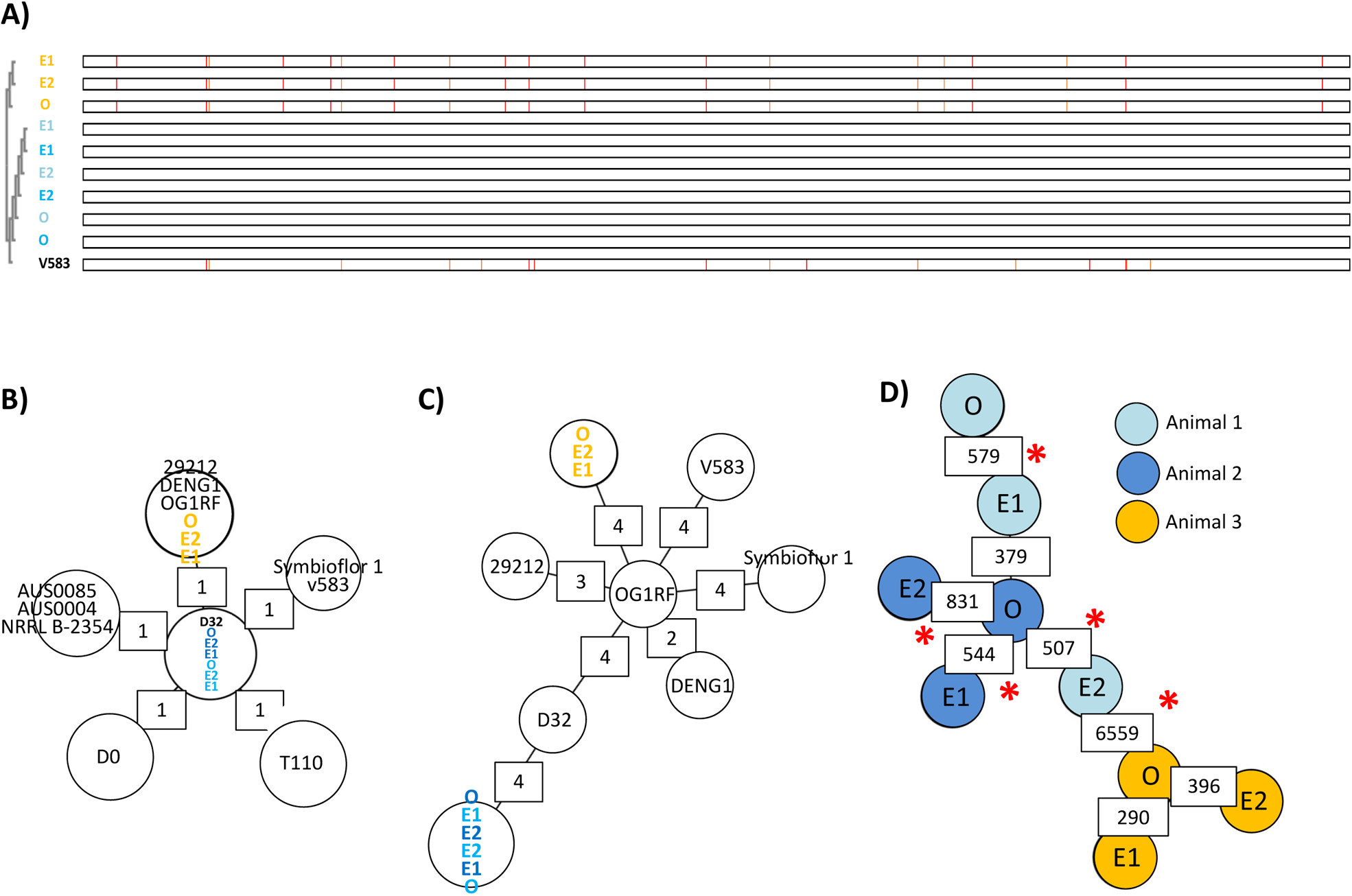
MSA, MST and cgMLST define two novel *E. faecalis* sequence types isolated across *N. nigricollis* venom and oral cavities. Blinded MSA (A) of the *KatA* gene sequence across the nine *E. faecalis isolates* obtained from *N. nigricollis* oral swabs (O) and two consecutive envenomation samples (E1 and E2) from three independent animals (light blue (N.nig01), dark blue (N.nig02) and orange (N.nig03)) defines two alleles distinct to the V583 reference sequence (bottom lane). Base conservation is defined by similarity to the animal 1 and 2 *KatA* sequence using BoxShade v.3.3.1 on mobyle.pasteur.fr. Each pixel column represents a different nucleotide with orange and red columns indicating increasingly different nucleotides. Blinded MST analysis of these nine isolates against B) *E. faecalis* and *E. faecium* reference genomes (distance calculations based on *E. faecium* MLST), C) *E. faecalis* reference genomes with partial incidence locus data removed, and D) a custom cgMLST schema derived from *E. faecalis* OG1RF, D32 and DENG1 including loci with partial data between all study isolates (8101 targets). Allelic differences in excess of 5% of the cgMLST schema are highlighted by ‘*’. Reference genomes: *E. faecalis*: V583, OG1RF, D32, DENG1, 29212, Symbioflor 1; *E. faecium*: T110; AUS0085, Aus0004, NRRL B-2354, DO.

**Figure S6:**
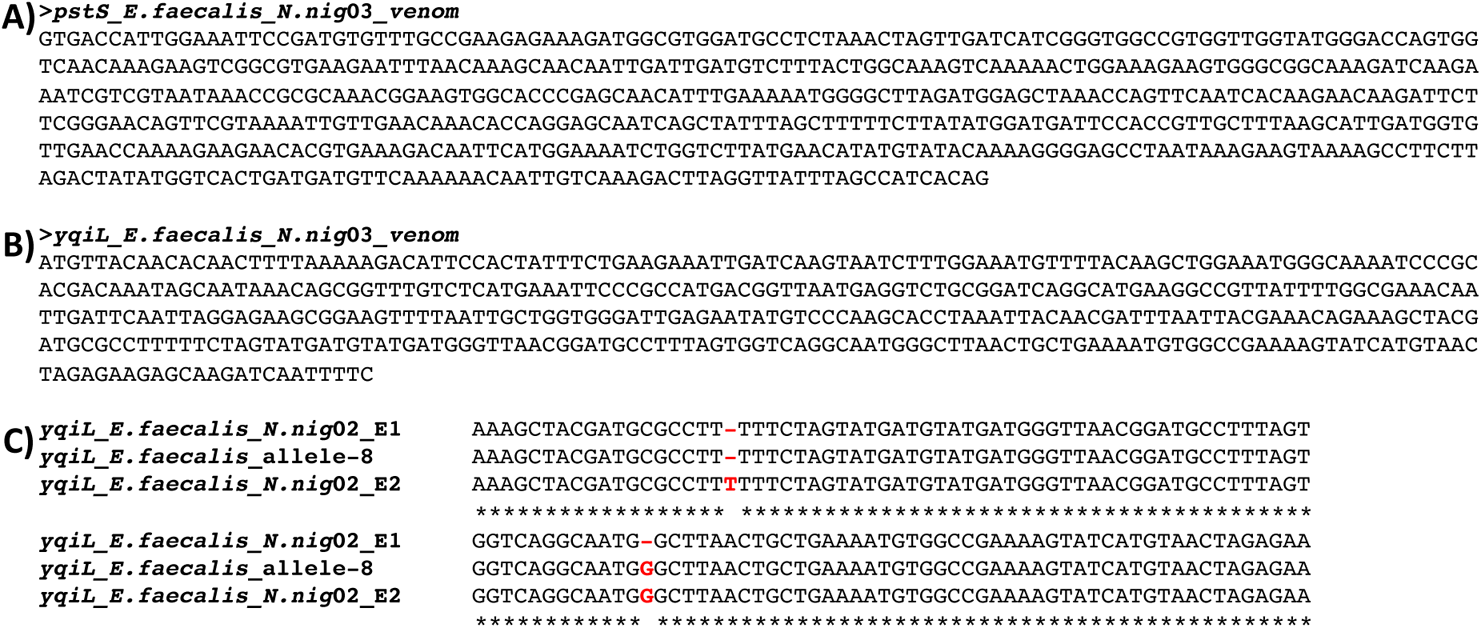
Novel *pstS* and *yqiL* allele sequences obtained from *N. nigricollis* venom-derived *E. faecalis.* The sequences of the novel *pstS* (A) and *yqiL* (B) alleles found in a novel *E. faecalis* sequence type obtained from *N. nigricollis* venom (animal 3). Clustal omega alignments of the *yqiL* sequences (C) from *E. faecalis* isolates derived from animal 2 venom against *E. faecalis yqiL* allele 8 found in the orally-derived isolate. The alignment is focused to positions 301-319 of the 436 nt allele and single base pair indels are highlighted in red.

**Figure S7:**
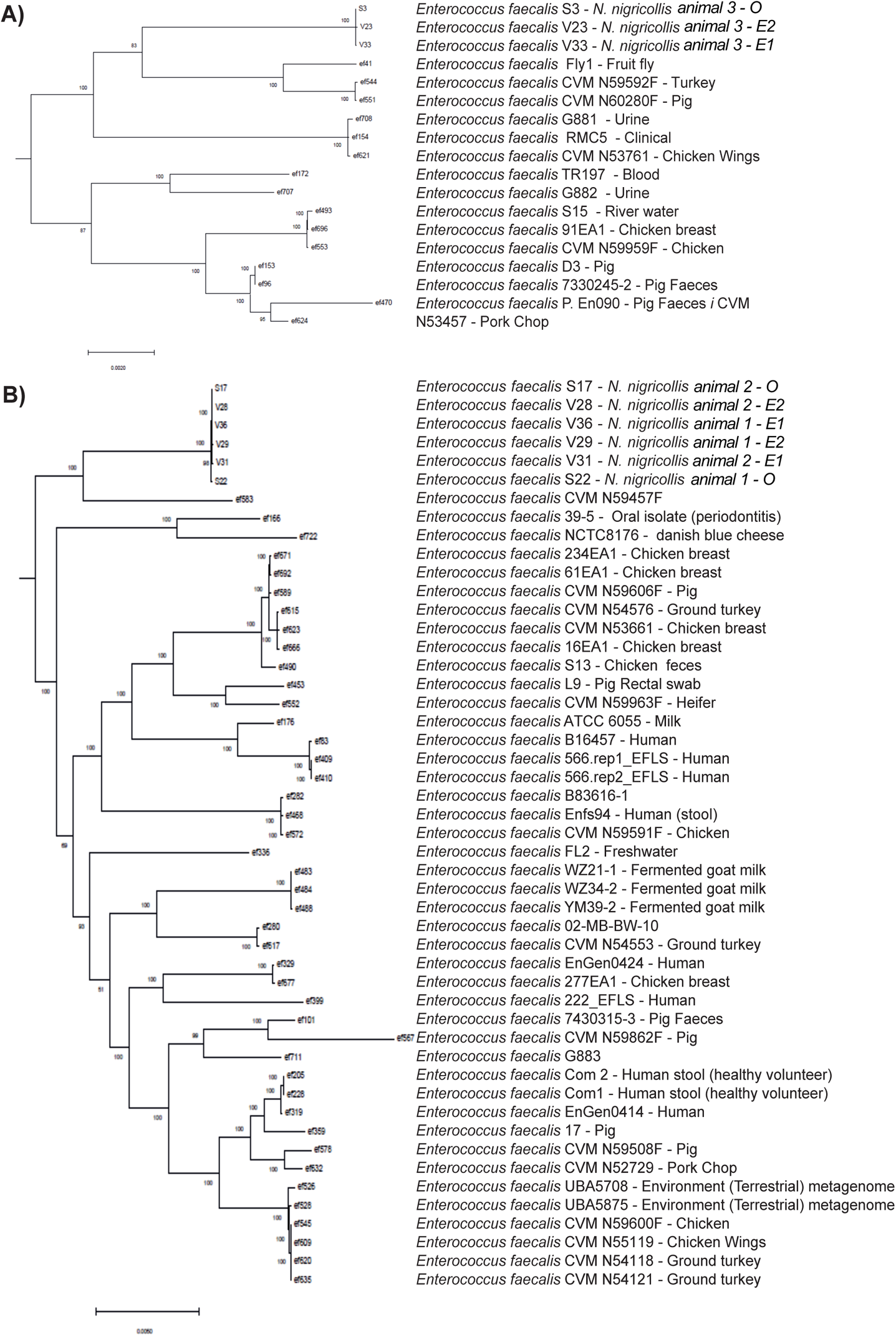
Source of *E. faecalis* strains with genomes closely related to venom-tolerant strains isolated from *N. nigricollis* venom. The genome record metadata available for the closest *E. faecalis* isolates related to (A) group A and (B) group B *N. nigricollis* venom isolates (subtrees extracted from the original 734 *E. faecalis* strain core genome tree) are depicted.

**Figure S8:**
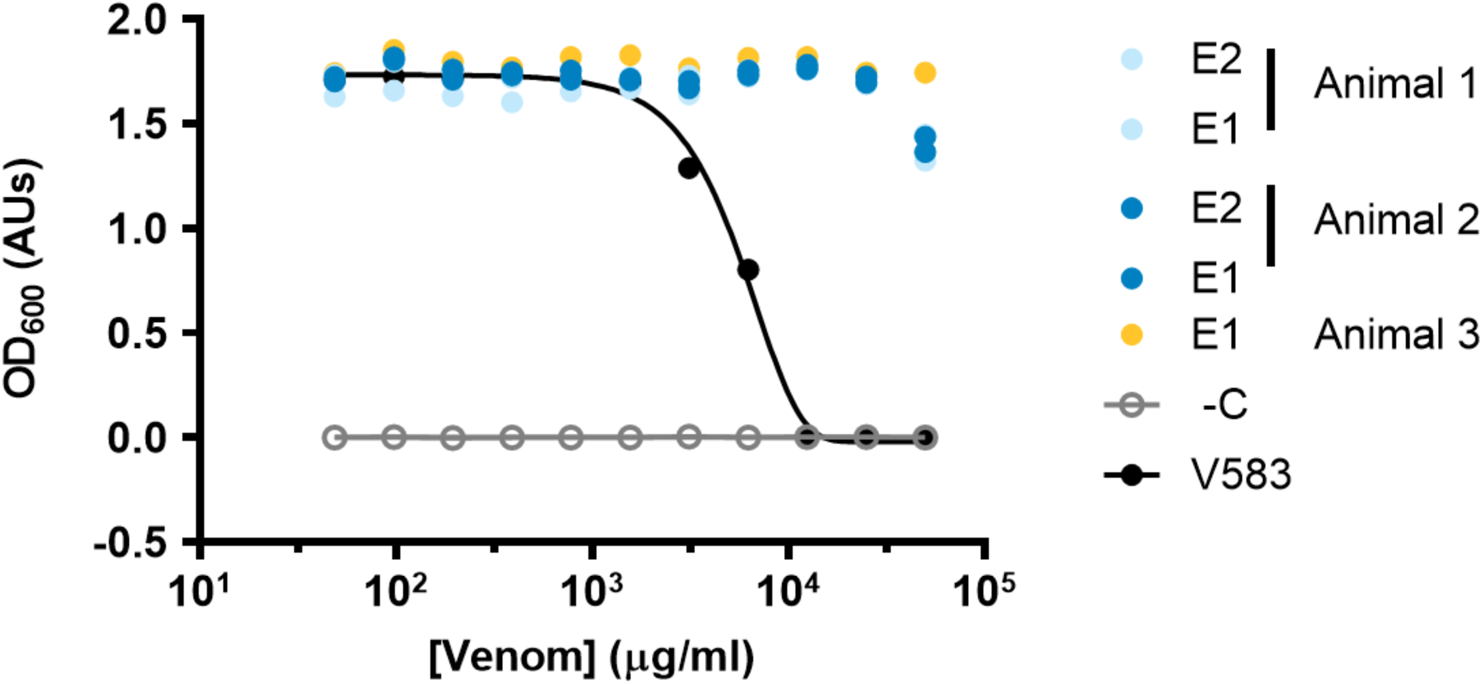
*N. nigricollis* venom*-*derived *E. faecalis* isolates resist the inhibitory effects of *N. nigricolis* lyophilised venom. The growth inhibitory effect of pooled, filter-sterilised, lyophilised *N. nigricollis* venom dissolved in brain heart infusion broth across 2-fold serial dilutions of 50 mg/ml was assessed for five *E. faecalis* isolates derived from *N. nigricollis* venom and the reference isolate V583 after 24 hr shaken incubation at 37°C by turbidity assessment at 600 nm. Data representative of 3 independent replicate experiments.

**Figure S9:**
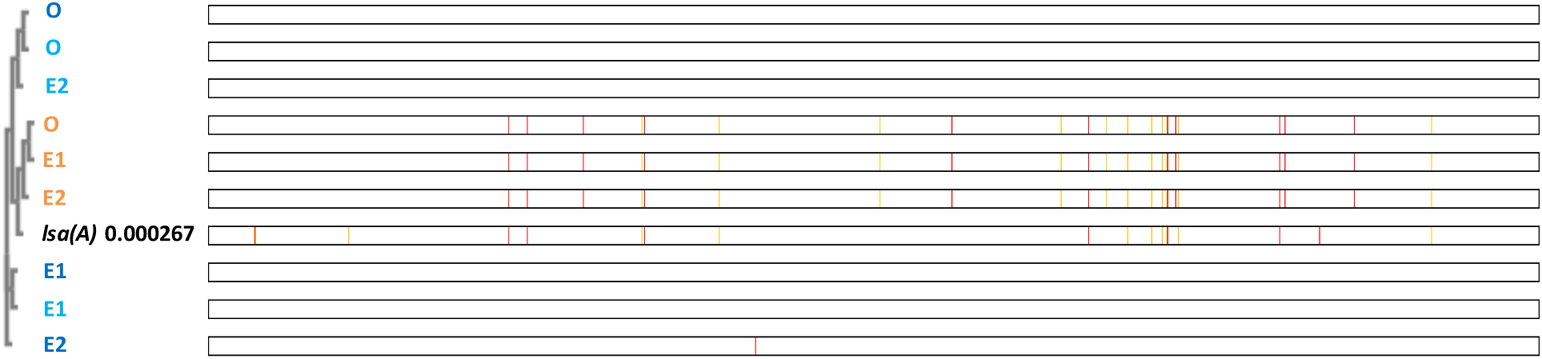
Multiple sequence alignment of the *lsa(A)* gene reinforces the clustering of the *N. nigricollis-*derived *E. faecalis* isolates. Blinded MSA of the *lsa(A)* antibiotic resistance gene sequence across the nine *E. faecalis isolates* obtained from *N. nigricollis* oral swabs (O) and two consecutive envenomation samples (E1 and E2) from three independent animals (light blue (N.nig01), dark blue (N.nig02) and orange (N.nig03)) defines two alleles distinct to the TX0263 reference strain gene sequence (accession no. AY737526.1). Base conservation is defined by similarity to the animal 1 and 2 *lsa(A)* sequence using BoxShade v.3.3.1 on mobyle.pasteur.fr. Each pixel column represents a different nucleotide with orange and red columns indicating increasingly different nucleotides.

**Table S2:** Isolate identification using BioMerieux biochemical strips.

| Animal no. | Sample type | Species (% Confidence Interval) |  |  |
| --- | --- | --- | --- | --- |
|  |  | 1 | 2 | 3 |
| 1 | O | *S. haemolyticus*<br>(47.9) | *S. aureus*<br>(17.9) | *S. saprophyticus*<br>(15.2) |
|  | E1 | *S. haemolyticus*<br>(35.8) | *S. warneri*<br>(32.0) | *S. homini*<br>(16.1) |
|  | E2 | *S. haemolyticus*<br>(35.8) | *S. warneri* (32.0) | *S. homini*<br>(16.1) |
| 2 | O | *S. haemolyticus*<br>(47.9) | *S. aureus*<br>(17.9) | *S. saprophyticus*<br>(15.2) |
|  | E1 | *P. vulgaris*<br>(62.7) | *P. penneri*<br>(35.2) | *P. mirabilis*<br>(2.0) |
|  | E1 | *S. haemolyticus*<br>(35.8) | *S. warneri*<br>(32.0) | *S. homini*<br>(16.1) |
|  | E2 | *S. haemolyticus*<br>(36.2) | *S. saprophyticus*<br>(19.5) | *S. aureus*<br>(18.2) |
| 3 | O | *S. cohnii ssp cohnii*<br>(35.5) | *S. warnerii*<br>(19.7) | *S. capitis*<br>(15.5) |
|  | E1 | *S. haemolyticus*<br>(35.8) | *S. saprophyticus*<br>(32.0) | *S. homini*<br>(16.1) |
|  | E2 | *S. haemolyticus*<br>(36.2) | *S. saprophyticus*<br>(19.5) | *S. aureus*<br>(18.2) |

O: oral swab sample
E1: envenomation 1
E2: envenomation 2

**Table S4:** Comparison of pTEF plasmid genomic elements in venom-resistant *E. faecalis* genomes groups isolates by animal of origin.

| Animal no. | Isolate origin | Average base reads per plasmid |  |  | Average base read ratio |  |  |
| --- | --- | --- | --- | --- | --- | --- | --- |
|  |  | pTEF1 | pTEF2 | pTEF3 | pTEF1:pTEF2 | pTEF2:pTEF3 | pTEF1:pTEF3 |
| 1 | O | 5.245 | 6.524 | 4.031 | 0.80 | 1.62 | 1.30 |
|  | E1 | 16.007 | 19.556 | 12.68 | 0.82 | 1.54 | 1.26 |
|  | E2 | 9.726 | 9.086 | 5.896 | 1.07 | 1.54 | 1.65 |
| 2 | O | 15.196 | 17.444 | 10.808 | 0.87 | 1.61 | 1.41 |
|  | E1 | 5.081 | 6.617 | 4.29 | 0.77 | 1.54 | 1.18 |
|  | E2 | 4.407 | 5.483 | 3.504 | 0.80 | 1.56 | 1.26 |
| 3 | O | 1.472 | 5.738 | 8.639 | 0.26 | 0.66 | 0.17 |
|  | E1 | 1.643 | 6.179 | 8.908 | 0.27 | 0.69 | 0.18 |
|  | E2 | 0.388 | 1.604 | 2.321 | 0.24 | 0.69 | 0.17 |

O: oral sample
E1: envenomation 1
E2: envenomation 2

**Table S10:**
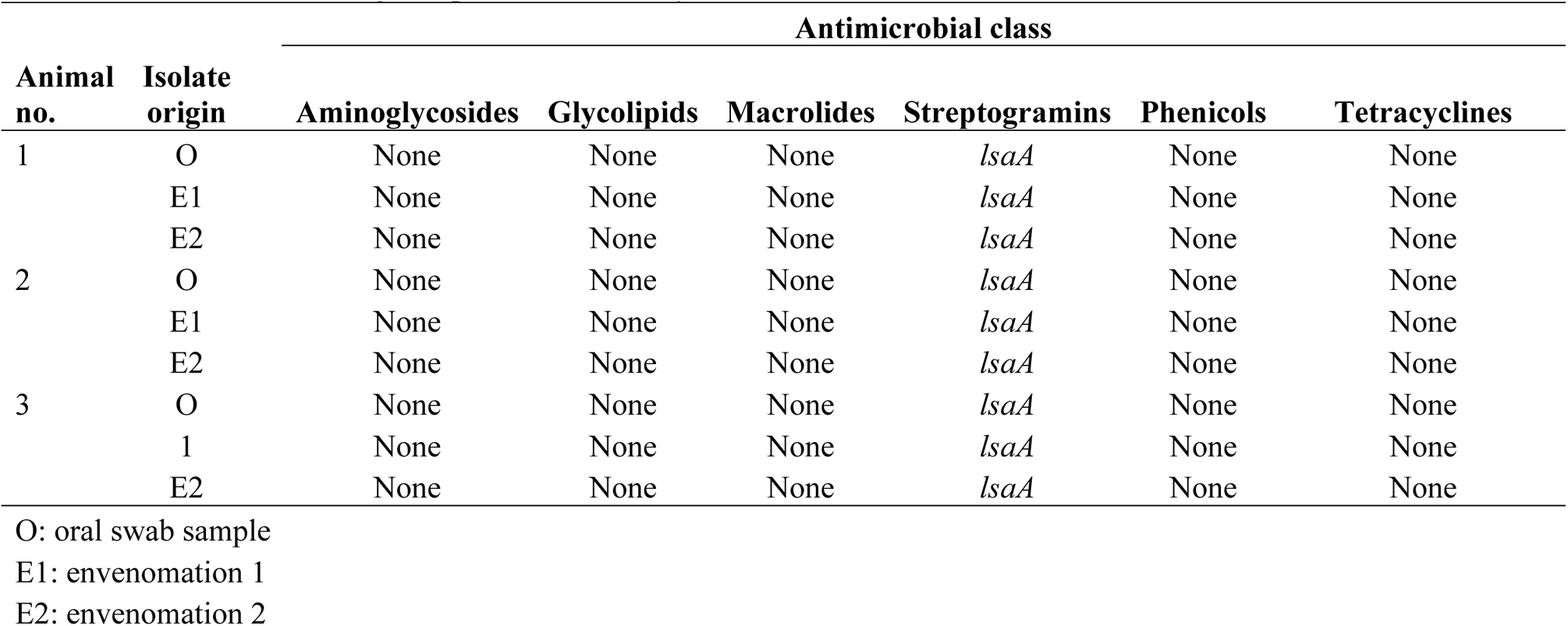
List of resistance genes present in each *E. faecalis* isolate for select antimicrobial classes.

| Animal no. | Isolate origin | Antimicrobial class |  |  |  |  |  |
| --- | --- | --- | --- | --- | --- | --- | --- |
|  |  | Aminoglycosides | Glycolipids | Macrolides | Streptogramins | Phenicols | Tetracyclines |
| 1 | O | None | None | None | *lsaA* | None | None |
|  | E1 | None | None | None | *lsaA* | None | None |
|  | E2 | None | None | None | *lsaA* | None | None |
| 2 | O | None | None | None | *lsaA* | None | None |
|  | E1 | None | None | None | *lsaA* | None | None |
|  | E2 | None | None | None | *lsaA* | None | None |
| 3 | O | None | None | None | *lsaA* | None | None |
|  | 1 | None | None | None | *lsaA* | None | None |
|  | E2 | None | None | None | *lsaA* | None | None |

O: oral swab sample
E1: envenomation 1
E2: envenomation 2

